# Associations Between Habitat Quality And Body Size In The Carpathian Land Snail *Vestia turgida*: Species Distribution Model Selection And Assessment Of Performance

**DOI:** 10.1101/2020.05.09.085746

**Authors:** V. Tytar, O. Baidashnikov

## Abstract

Species distribution models (SDMs) are generally thought to be good indicators of habitat suitability, and thus of species’ performance, consequently SDMs can be validated by checking whether the areas projected to have the greatest habitat quality are occupied by individuals or populations with higher than average fitness. We hypothesized a positive and statistically significant relationship between observed in the field body size of the snail *V. turgida* and modelled habitat suitability, tested this relationship with linear mixed models, and found that indeed, larger individuals tend to occupy high-quality areas, as predicted by the SDMs. However, by testing several SDM algorithms, we found varied levels of performance in terms of expounding this relationship. Marginal R^2^, expressing the variance explained by the fixed terms in the regression models, was adopted as a measure of functional accuracy, and used to rank the SDMs accordingly. In this respect, the Bayesian additive regression trees (BART) algorithm (Carlson, 2020) gave the best result, despite the low AUC and TSS. By restricting our analysis to the BART algorithm only, a variety of sets of environmental variables commonly or less used in the construction of SDMs were explored and tested according to their functional accuracy. In this respect, the SDM produced using the ENVIREM data set (Title, Bemmels, 2018) gave the best result.

## 1. Introduction

Information on where species occur underlies nearly every aspect of managing biodiversity (Franklin 2010), but knowledge of distributions is often coarse or incomplete. Species distribution models, SDMs (closely related to ecological niche models, ENMs, bioclimate-envelope modelling etc.) provide a tool used to derive spatially explicit predictions of environmental suitability for species (Guisan et al., 2017) by employing suitability indices. Suitability indices describe the relationship between habitat suitability score and a given environmental variable of a target species. Habitat suitability is a way to predict the suitability of habitat at a certain location for a given species or group of species based on their observed affinity for particular environmental conditions (Yi et al., 2016; Ma, Sun, 2018). SDMs therefore have been widely used for predicting distributions of species in terrestrial, freshwater and marine environments, and across taxa from many biological groups (Elith, Leathwick, 2009), with increasing numbers of publications each year (Robinson et al., 2011; Brotons, 2014). SDMs have shown to be efficient in biodiversity research considering climate changes (Barbet-Massin et al., 2011; Visconti et al., 2016), conservation planning (Kremen et al., 2008), invasive species and pest risk assessments (Gallien et al., 2012; Jeger et al., 2018), pathogen spread (Schatz et al., 2017), rewilding projects (Jarvie, Svenning, 2018), and a huge number of other issues ranging from mapping snake bite risk (Yañez-Arenas et al., 2016) to Pygmy presence in Central Africa (Olivero et al., 2016).

SDM tools generally correlate species’ occurrence patterns with environmental variables, which are frequently selected from an array of ‘bioclimatic’ indices (Hijmans et al., 2005; Kriticos et al., 2012; etc.) and thus focus on the abiotic conditions affecting species distributions (Busby, 1991). Recently, more studies include biotic covariates (Wisz et al., 2013; Si-Moussi et al., 2019), motivated by the need to account for more directly explanatory variables and resources, although dependencies between species (for instance, competition) may be correlated indirectly through latent abiotic variables. So in general, keeping in mind the geographic scale, adaptation to abiotic factors allows to assume adaptation to biotic interactions too. For example, temperature comprises a large set of ecophysiologically relevant variables (Dahl, 1998), but even simple temperature variables, like annual mean temperature, covary spatially with many broad-scale biotic patterns at regional and global scales (Leith, Whittaker, 1975). Apparently the success of bioclimate-envelope modelling comes from this strong spatial covariance between easily measured abiotic variables and the poorly understood and largely unknown ecologically critical biotic variables (Jackson et al., 2009).

Traditionally, determining environmental and climatic features that characterize the species’ niche and are responsible for shaping their distribution would require laborious field measurements of key environmental variables in natural populations (Nakazato et al., 2010; Warren et al., 2020). Importantly, the use of SDMs has allowed to identify such driving factors, but SDM construction involves many decisions which may adversely affect model predictions, including the choice of modelling algorithms (Warren et al., 2020). Choices regarding optimal models and methods are typically made based on discrimination accuracy, which only measures whether a model assigns higher suitability values to presence points than it does to background or absence points (Gurgel-Gonçalves et al., 2012), and conclusions have been made of the inability of current evaluation metrics to assess the biological significance of distribution models (Fourcade et al., 2018).

Predictions from SDMs are generally thought to be good indicators of habitat suitability, and thus of species’ performance. An implicit assumption of the SDMs is that the predicted ecological niche of a species actually reflects the adaptive landscape of the species, so in sites predicted to be highly suitable, species would have maximum fitness compared to in sites predicted to be poorly suitable (Zizhen, Hong, 1997; Nagaraju et al., 2013). Therefore these models potentially can be validated by checking whether the areas projected to have the greatest habitat quality are occupied by individuals or populations with higher than average fitness (Mammola et al., 2019), in other words check the SDMs functional accuracy. For instance, a positive correlation (r = 0.5) was found between the growth rate of a wild grass carp, *Ctenopharyngodon idella* (Valenciennes, 1844), and habitat quality for the species as projected by a maximum entropy model (Wittmann et al., 2016). In another case modelled habitat quality was positively associated with maximum body and egg case size in a spider species, Vesubia jugorum (Simon, 1881) (Mammola et al., 2019). Yet, in the few studies that have explicitly tested the relationship between habitat quality and species traits, not always such relationship was found (Mammola et al., 2019). The problem could be that it is often not clear which measurable biological phenomena should be correlated with suitability estimates from SDMs, moreover when many of the measurable phenomena that are potentially related to suitability have not been quantified in detail and as such are merely unavailable for model validation (Warren et al., 2020).

In this study we attempted to highlight important variables shaping the current niche of a terrestrial gastropod, *Vestia turgida* (Rossmassler, 1836), found primarily in the Carpathian Mountains, with a focus on the relationships between habitat quality and species traits, consequences these may have in terms of model selection and performance. We argue that the habitat suitability of a species as predicted by the ecological niche model may also reflect the adaptive landscape of the species. Indeed, species should have a higher performance in the core of their niche (i.e. where conditions are more suitable) than at their edges (Pulliam, 2000).

- First, using the existing natural distribution data of the species and a variety of environmental variables, we generate ecological niche model predictions on the habitat suitability of *V. turgida* in the Ukrainian Carpathians by employing a number of algorithms commonly used or recently developed for constructing SDMs.
- Second, we evaluate the model predictions both in terms of discrimination accuracy, using conventional criteria, and functional accuracy by using body size as a measure of fitness and testing whether the degree of predicted habitat suitability correlates positively with observations of body size. The null hypothesis was that no correlation exists between observed body size and modelled habitat suitability, r = 0. We hypothesized a positive, significant relationship.
- Finally we rank the SDM outputs and select the ‘best’ modelling approach to analyze the environmental niche of the species to see which of the employed sets of environmental variables promote better performance of the SDMs in terms of functional accuracy.

## 2. Species and study area

*V. turgida* is a species of air-breathing land snail, a terrestrial pulmonate gastropod mollusk in the family *Clausiliidae*, the door snails, all of which have a clausilium, a roughly spoon-shaped “door”, which can slide down to close the aperture of the shell (Лихарев, 1969). The species is considered an endemic Carpathian snail, though sporadically met in the Dnestr Basin of Podolia in Ukraine. It is widely distributed in the Carpathian Region with numerous localities especially in Slovakia, Poland and Ukraine. Several isolated populations are threatened, especially due to changes in forest management and water drainage. However, the whole species is considered of Least Concern (LC) (Walther, 2017). Nevertheless, isolated relict subpopulations far off the main range as well as marginal populations could be endangered by human encroachment and climate change.

According to Kerney et al. (1983), *V. turgida* occurs in very moist woodland, under logs and ground litter. It is supposed that the litter and bacteria decomposing dead wood are the main components of the diet of clausiliids (Fog, 1979). The species ascends to 2100 m in the Carpathians (the Tatra Mountains) (Dyduch-Falniowska, 1991), whereas in the Podolia snails are found at a much lower height (down to 200 m of even less).

The study area largely encompasses the range of the Ukrainian Carpathians (48°32 N, 23°38 E), which extends over an area of about 24 000 sq. km. The study area lies at an altitude of 95–2030 m, although 94% of the mountains are < 1200 m. The highest elevations are located in the southern parts of the Ukrainian Carpathians, while the south-west (bordering Romania), west (bordering the Transcarpathian Lowland of Ukraine) and north-west (bordering Poland)Carpathians are characterized by extensive valley systems and relatively gentle slopes. Precipitation of 500–1400 mm/year feeds a dense network of rivers (Голубець, 1988). The July (warmest month) temperature varies from 20°C at the southern edge of the Carpathians and 18°C in the north to 6°C on the highest peaks (Геренчук, 1968; Kuemmerle et al. 2009). Winter temperatures range from −3°C to −10°C. The mountains are dominated by *Fagus sylvatica* and *Picea abies* forests, replaced by *Pinus mugo* and *Juniperus communis* in the subalpine and grasslands in the alpine belts (Геренчук 1968; Kuemmerle et al. 2009).

Next to this area, were the species is more or less sporadically found, is the Dnister basin of Podolia, covering around 24 500 sq. km, with an average altitude of 320-350 m. The climate is temperate-subcontinental with a mean annual temperature of about 7–9°C and 600–650 mm annual precipitation (Клімат України, 2003). The relief of the area is dissected by numerous river valleys into distinct ridges. About 10-15% of the area is occupied by the forest vegetation comprised of oak-hornbeam-beech stands.

### 2.2 Species distribution modelling

#### 2.2.1 Collection data

In 1985–1986, 1989–1992 and 2004 snails were collected by hand at 94 georeferenced stations sites at elevations up to 1527 m a.s.l. Sampling intensity varied over geography, therefore to minimize spatial sampling heterogeneity, we aggregated data at the resolution of the environmental predictors to avoid inflation of the number of presences.

With a LOMO Binocular Stereo Microscope MBS-1, we measured in 1 016 specimens of *V. turgida* three morphological traits related to body size: shell height (H), shell diameter (D) and number of whorls (Wh) using a conventional standard (Лихарев, Раммельмейер, 1952).

#### 2.2.2 Environmental predictors

In most cases environmental predictors are selected based on the availability and experience that the variables show correlation with the species distribution (Guisan, Zimmerman, 2000). Because of the habitat complexity it is difficult to single out which factors play a crucial role in controlling molluscs distribution (Sulikowska-Drozd, 2005), but for the majority of terrestrial gastropods their occurrences are considered to be determined by several factors, such as pH and calcium content (Nekola, Smith, 1999; Martin, Sommer, 2004), drainage (Paul, 1978), altitude (Cowie et al., 1995), shelter possibilities (South, 1965), humidity (Martin, Sommer, 2004), plant composition, and plant diversity (Barker, Mayhill, 1999). Important environmental factors emerging from these studies are moisture conditions, vegetation structure and soil pH, which is related to soil calcium content (Astor, 2014).

Under these circumstances we might expect temperature and precipitation variables, together with their various combinations, to be important (Hof, 2011). Climate variables used in SDMs are assumed to reflect the physiological constraints on the study species that affect where they can survive in the wild (Kearney, Porter, 2009), although many commonly used SDM variables have been shown to often neglect important physiological factors (Gardner et al., 2019). Nevertheless, we employ climate variables anticipating their wider impacts, by being closely linked to the energy available in the ecosystem or the length of the growing seasons, plant growth, species’ spatial variation patterns owing to moisture availability, operating through variations in plant productivity, impact on soil properties, etc.

We used the widely accepted bioclimatic potential predictor variables for species distribution and suitability analysis (Hijmans et al., 2005). These bioclimatic predictors are ecologically more sensitive to differentiate the physio-ecological tolerances of a habitat (Thompson et al., 2009) than simple temperature and precipitation predictors (Graham, Hijmans, 2006; Kumar et al., 2009). Information on the bioclimatic parameters was collected as raster layers from the WorldClim website (http://www.worldclim.org/current) with a spatial resolution of 30 arc seconds. These variables indicate a general trend of precipitation and temperature, extremity and seasonality of temperature.

Former studies have shown a strong influence of topography on both biotic and abiotic factors in study areas (Homeier et al. 2010; Werner et al. 2012; Svenning et al. 2009) and topography variables are observed to make an extremely high (up to 90%) contribution to species distribution models (Dudov, 2017). In this study, topographic variables (e.g. elevation, slope, aspect etc.) are used as proxies for environmental factors such as insolation, wind exposure, hydrological processes etc., affecting the quality of the species’ habitat. Topographical variables were based on the SRTM data set that is available at http://srtm.csi.cgiar.org. Derived topographic variables were calculated using the open source software SAGA GIS (Conrad et al., 2015).

Valuable remotely sensed predictors for site quality and forest species communities also include vegetation indices such as the normalized difference vegetation index (NDVI), which has been widely used as surrogate of primary productivity and vegetation density (Pettorelli et al., 2005). Vegetation data include maps NDVI obtained from satellite images by NASA and processed at Clark Lab (www.clarklabs.org). Means and deviations were computed over an 18-year period (from 1982 to 2000) and original NDVI real values (from −1 to +1) were rescaled to a range from 1 to 255 (byte format).

Considering that vegetation is highly influenced by edaphic variables, we also examined soil properties. In many studies on land snails, particular attention was paid to soil chemical parameters, as snails have a high demand of calcium for shell formation (Martin, Sommer, 2004). However, several studies confirm the importance of a range of soil characteristics as determinants of gastropod distribution (Ondina et al., 2004). Soil properties, including physical and chemical features, were downloaded from SoilGrids (www.soilgrids.org), a system for global digital soil mapping (Hengl et al., 2014).

In this study we used for modelling purposes a recently reconsidered in terms of biological significance set of 16 climatic and two topographic variables (the ENVIREM dataset, downloaded from http://envirem.github.io), many of which are likely to have direct relevance to ecological or physiological processes determining species distributions (Title, Bemmels, 2018). These variables are worth consideration in species distribution modeling applications, especially as many of the variables (in particular, potential evapotranspiration) have direct links to processes important for species ecology.

#### 2.2.3 Calibration area

We calibrated and projected SDMs within the spatial extent of the Ukrainian Carpathians. Because true absence data is not available, pseudo-absence data was generated in locations with contrasting environmental conditions (Barbet-Massin et al., 2012), using the BCCVL application (Hallgren et al., 2016).

#### 2.2.4 Modelling methods

There exists a large suite of algorithms for modelling the distribution of species, but because there is no single ‘best’ algorithm some authors have reasonably concluded that niche or distribution modelling studies should begin by testing a suite of algorithms for predictive ability under the particular circumstances of the study and choose an algorithm for a particular challenge based on the results of those tests (Qiao et al., 2015).

Accordingly, we assessed the relative performance of various categories of SDM algorithms: BIOCLIM (Busby, 1991; Booth et al., 2014), Generalized Linear Models (GLMs, Guisan et al., 2002), MaxLike (Royle, J.A. et al., 2012), Random forests (Breiman, 2001), Boosted Regression Trees (Elith et al., 2008), Support Vector Machines (SVMs; Vapnik, 1998), and Bayesian additive regression trees (BART, Carlson, 2020).

SDM methods, excluding BART, were employed using the “sdm” package within the statistical software R (Naimi, Araújo, 2016), following the recommended by the authors default settings. Models were evaluated by 10-fold cross-validation using 30% of the occurrence dataset, and incorporating the aforementioned pseudo-absence data.

Initially we fitted models that included a selection of non-collinear environmental variables from the entire set based on the variance inflation factor (VIF, Marquardt, 1970): strongly collinear variables (VIF>10) were discarded. Subsequently automated variable set reduction was employed.

In terms of discrimination accuracy model performance was evaluated using two commonly used validation indices: the area under a receiver operating characteristic (ROC) curve, abbreviated as AUC, and the True Skill Statistic (TSS). The AUC validation statistic is a commonly used threshold independent accuracy index that ranges from 0.5 (not different from a randomly selected predictive distribution) to 1 (with perfect predictive ability). Models having AUC values >0.9 are considered to have very good, >0.8 good and >0.7 useful discrimination abilities (Metz, 1978). The TSS statistic ranges from −1 to +1 and tests the agreement between the expected and observed distribution, and whether that outcome would be predicted under chance alone (Allouche et al., 2006; Liu et al., 2009). A TSS value of +1 is considered perfect agreement between the observed and expected distributions, whereas a value <0 defines a model which has a predictive performance no better than random (Allouche et al. 2006). TSS has been shown to produce the most accurate predictions (Jiménez-Valverde et al., 2004). Values of TSS < 0.2 can be considered as poor, 0.2–0.6 as fair to moderate and >0.6 as good (Landis, Koch, 1977)

#### 2.2.5 Relationships between body size and habitat quality

Geographic variation in size has been found to be correlated with a variety of abiotic and biotic environmental factors. For instance, shells in the land snail *Albinaria idaea* (*Gastropoda: Clausiliidae*) are larger in regions of high temperature, and are generally larger in areas with higher rainfall (Welter-Schultes, 2000). In an extensive literature review shell size in terrestrial gastropods individualistic responses have been noted along moisture, temperature/insolation, and calcium availability gradients (Goodfriend, 1986), although the author could not identify universal ecological predictors. Most likely synergetic interactions between them could be the best explanation of the size variations resulting from the influence of local environmental and/or climate factors (Proćków et al., 2017), where maximum sizes are attained at environmental optima (Rensch, 1932, 1939; Терентьев, 1970). Our assumption is that these conditions are adequately reflected in the projected habitat quality for the species.

There was a high degree of correlation among the shell traits related to body size (Pearson correlations between shell height (H) and the shell diameter (D), and the number of whorls (Wh) was 0.90 and 0.85, respectively. Therefore shell height (H) was selected as a representative proxy of body size.

We tested the relationship between body size and projected habitat quality with linear mixed models (LMMs) that we fitted using the ‘Mixed Model’ module in the jamovi computer software (The jamovi project, 2020). This mixed method allowed to address the fact that because we measured multiple individuals from the same populations, we violated the models’ assumption of spatial independence. The sampling location was included as a random factor and the variance explained by the fixed terms in the regression models was expressed as marginal R^2^ and adopted as a measure of functional accuracy.

## 3 Results

### 3.1 Species distribution modelling and selection

After removing duplicate occurrences, we used 85 occurrences to generate the SDMs. We selected thirteen non-collinear variables for constructing the models. These represent the bioclimate (mean diurnal range, isothermality, precipitation seasonality, and precipitation of warmest quarter), topography (eastness, northness, slope, and topographic position index), NDVI for February, April and June, and soil properties (cation exchange capacity and silt content).

The outputs of the SDM algorithms varied in terms of discrimination accuracy evaluated by the AUC and TSS (Table 1).

**Table 1.** Discrimination accuracy of employed SDM algorithms*

| SDM methods | AUC | TSS |
| --- | --- | --- |
| BIOCLIM | 0.65 | 0.31 |
| Generalized Linear Model (GLM) | 0.91 | 0.75 |
| MaxLike | 0.88 | 0.70 |
| Random forests (RF) | 0.98 | 0.89 |
| Boosted Regression Trees (BRT) | 0.93 | 0.78 |
| Support Vector Machines (SVM) | 0.96 | 0.81 |
| Bayesian additive regression trees (BART) | 0.89 | 0.61 |

\*see abbreviations in the text

According to these results, the Random forests (RF) model demonstrates the best preformance (AUC = 0.98, TSS = 0.89), whereas the performance of the BIOCLIM model is behind the rest of the employed SDMs (AUC = 0.65, TSS = 0.31).

In all cases we found a positive, significant (p < 0.001) relationship between shell height (H) and habitat quality as projected by the models, with larger individuals in high-quality areas (Table 2).

**Table 2.** Functional accuracy of employed SDM algorithms*

| SDM methods | Estimated $\beta \pm SE$ | $R^2$ | AIC** |
| --- | --- | --- | --- |
| BIOCLIM | 6.02±1.734 | 0.107 | 4082.0 |
| Generalized Linear Model (GLM) | 3.33±0.580 | 0.098 | 3008.3 |
| MaxLike | 3.73±0.997 | 0.054 | 3016.6 |
| Random forests (RF) | 6.43±1.213 | 0.142 | 3013.0 |
| Boosted Regression Trees (BRT) | 9.48±1.376 | 0.192 | 2991.5 |
| Support Vector Machines (SVM) | 3.26±0.703 | 0.075 | 3008.1 |
| Bayesian additive regression trees (BART) | 5.88±0.845 | 0.210 | 2987.4 |

\*see abbreviations in the text
\*\* AIC – Akaike information criterion (Aho et al., 2014)

Based on these findings, the most biologically meaningful model has been constructed using the Bayesian additive regression trees (BART) algorithm, which has outperformed other SDM algorithms of the employed suite, with the highest marginal R^2^ (0.210) and lowest AIC (2987.4). The linear relationship, derived from the linear mixed model, between habitat quality predicted by the BART model and shell height is shown in Figure 1. Treading on the heels of the BART model and displaying good performance is the Boosted Regression Trees (BRT) model, with a marginal R^2^ only somewhat lower (0.192) and AIC slightly higher (2991.5). Interestingly, BIOCLIM, the vet of SDMs (Nix, 1986), according to the applied criteria (aside from AIC), appears to have performed better than some of the other algorithms in the suite, including machine learning methods.

**Figure 1.**
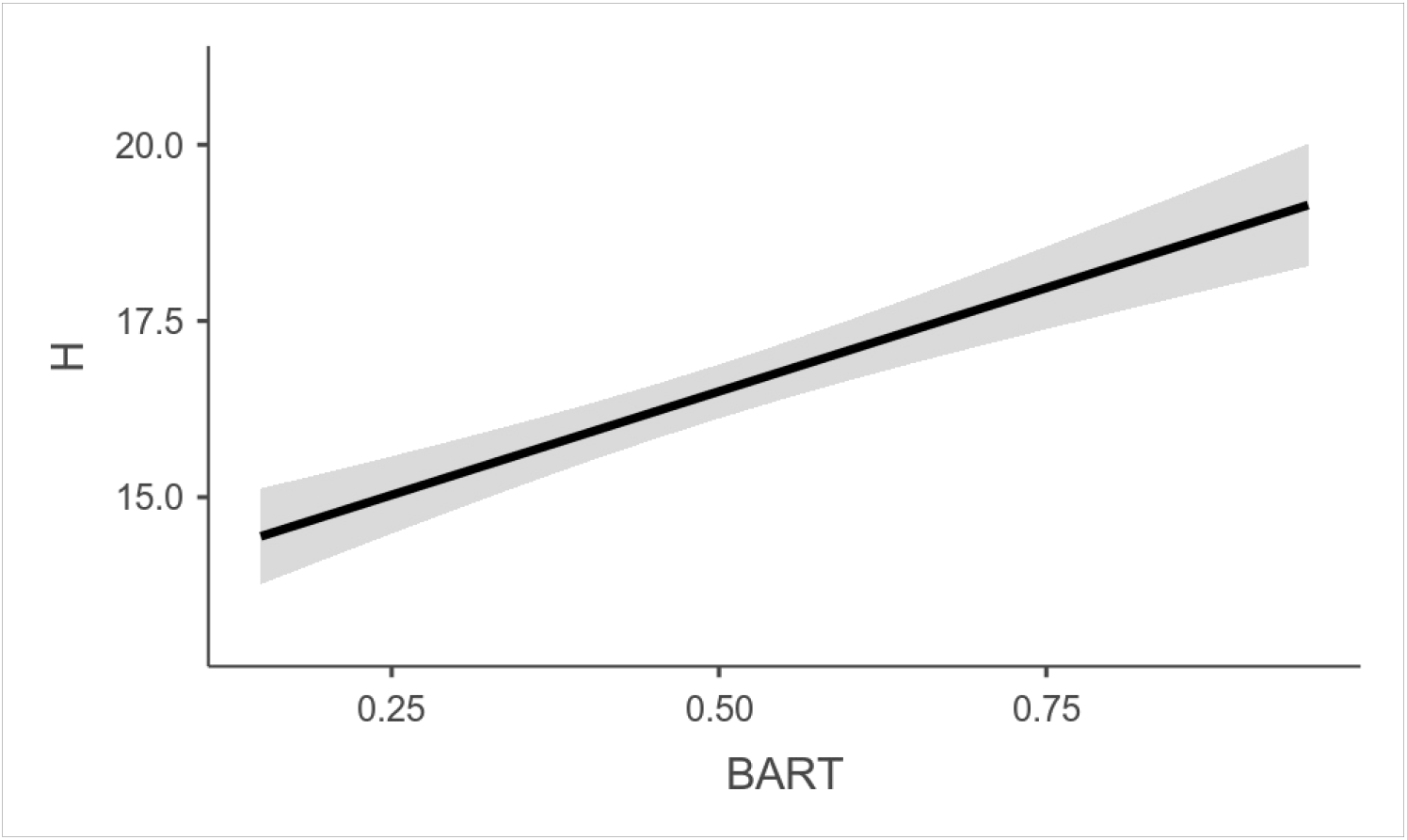
Linear relationship (solid line) and 95% confidence interval (gray area) between habitat quality predicted by the BART model (*x-axis*) and shell height (H in millimeters, *y-axis*), derived from the linear mixed model.

### 3.2 Analysis of the environmental niche using BARTs

Based on the results of the preceding tests, the Bayesian additive regression trees (BART) algorithm has been selected to perform an indepth analysis of the niche of the snail *V. turgida* in relation to environmental predictors listed in **2.2.2**.

Bayesian additive regression trees (BART) are a new alternative to other popular classification tree methods. In computer science, BARTs are used for everything from medical diagnostics to self-driving car algorithms, however they have yet to find widespread application in ecology and in predicting species distributions. Running SDMs with BARTs has recently been greatly facilitated by the development of an R package, ‘embarcadero’ (Carlson, 2020), including an automated variable selection procedure being highly effective at identifying informative subsets of predictors. Also the package includes methods for generating and plotting partial dependence curves.

#### 3.2.1 Bioclimatic variables

Nineteen bioclimatic variables from the WorldClim base were used in the species distribution modelling (their codes and names are available here: https://worldclim.org/data/bioclim.html; accessed 26.04.2020).

Five bioclimatic variables were identified as an informative subset of predictors: BIO17 = Precipitation of Driest Quarter, BIO16 = Precipitation of Wettest Quarter, BIO5 = Max Temperature of Warmest Month, BIO9 = Mean Temperature of Driest Quarter, and BIO10 = Mean Temperature of Warmest Quarter, which capture the basic bioclimatic requirements of the snail. On-topic accuracy measures for the model are presented in Table 3.

**Table 3.** Accuracy measures for the BART model based on bioclimatic variables

| AUC | TSS | Estimated $\beta \pm SE$ | t-test | p | R<sup>2</sup> | AIC |
| --- | --- | --- | --- | --- | --- | --- |
| 0.854 | 0.620 | 7.06±0.991 | 7.13 | <0.001 | 0.249 | 2968.8 |

Within this subset the modelling brought out the high importance of BIO17 = Precipitation of Driest Quarter. The driest quarter in the study area broadly coincides with the cold season, therefore BIO17 can be considered a proxy for snow depth. Snow is a highly effective insulator and can provide a significant buffer against winter temperature extremes (Sturm et al., 2001; Zhang, 2005; Nicolai, Ansart, 2017). Here, in the case of *V. turgida*, highly suitable areas are those where the cold season precipitation is above 170 mm (projected habitat suitability above 70%), whereas below that level suitability rapidly decreases to a projected 30% and less (Figure. 2).

**Figure 2.**
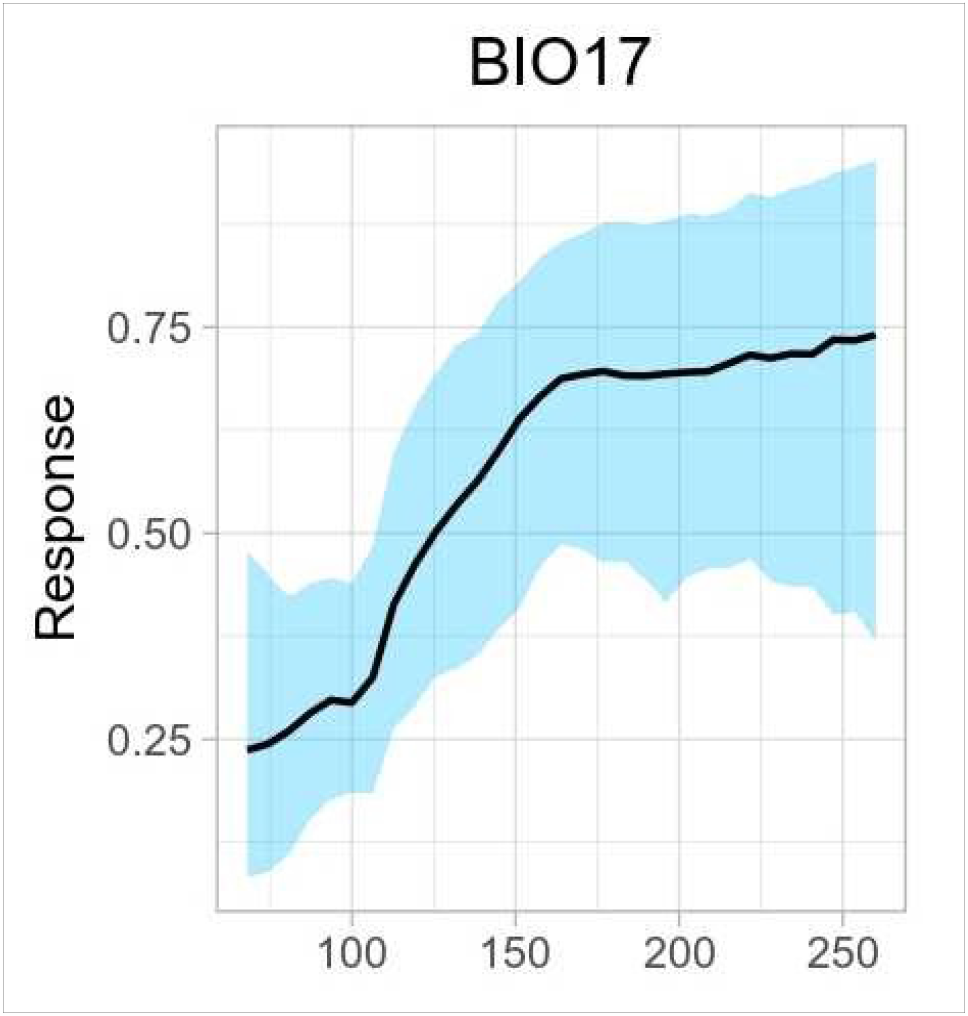
Partial dependence plot for BIO17 = Precipitation of Driest Quarter; blue area = 95% confidence interval

A positive, significant relationship was found between shell height and habitat quality as projected by the model, with larger individuals in bioclimatically more suitable areas.

#### 3.2.2 Topographic variables

We used a set of topographic variables including elevation (although there are opposing views on whether to include elevation as a predictor in SDMs or not; see, for example, Hof et al., 2012), slope, aspect (eastness, northness), terrain roughness index (Wilson et al., 2007) SAGA-GIS topographic wetness index (Boehner et al., 2002), and topographic position index (Guisan et al., 1999). Importantly, strong relationships between body size of *V. turgida* and elevational gradients have been reported (Байдашников, 1985; Sulikowska-Drozd, 2001).

Three topographic variables were identified as an informative subset of predictors: elevation, SAGA-GIS topographic wetness index (TWI), and terrain roughness index (tri). Corresponding accuracy measures for the model are presented in Table 4.

**Table 4.**
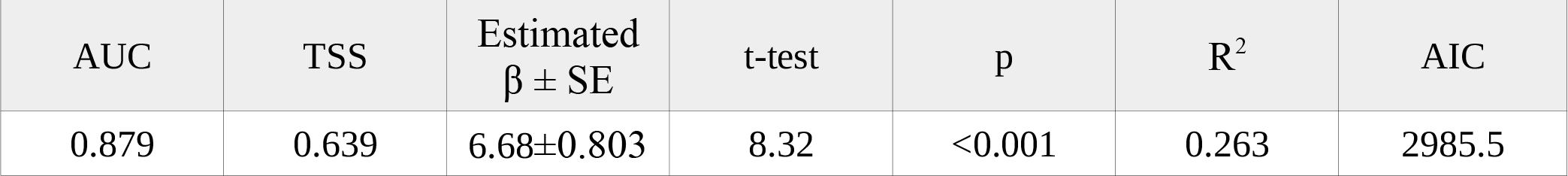
Accuracy measures for the BART model based on topographic variables

There is a general hump-shaped relationship between the habitat suitability values and elevation. Highest projected habitat suitability, using a 50% habitat suitability threshold (Waltari, Guralnick, 2009), is shown to occur between elevations of around 200 and 580 m a.s.l.

TWI, another topographic variable of recognized importance, calculates the capacity of water accumulation of each pixel in a watershed. Pixels with higher TWI values have higher capacity of water accumulation (Besnard et al., 2013) or, in other words, being “wetter”. The index is highly correlated with several soil attributes such as horizon depth, silt percentage, organic matter content, and phosphorus (Moore et al., 1993), can be used to simulate the status of soil moisture, which also has an influence on soil pH (Song, Cao, 2017). In our case, increasing values of TWI in relation to projected habitat suitability show a steady downward trend (Figure 3), meaning habitats that are “too wet” do not favour the species. Indeed, *V. turgida* occurs in very moist woodland (Kerney et al., 1983), however it has also been shown that the snail avoids very damp places (Urbański, 1939), so presumably our modelling results are consistent with these findings based upon observations made in the field.

**Figure 3.**
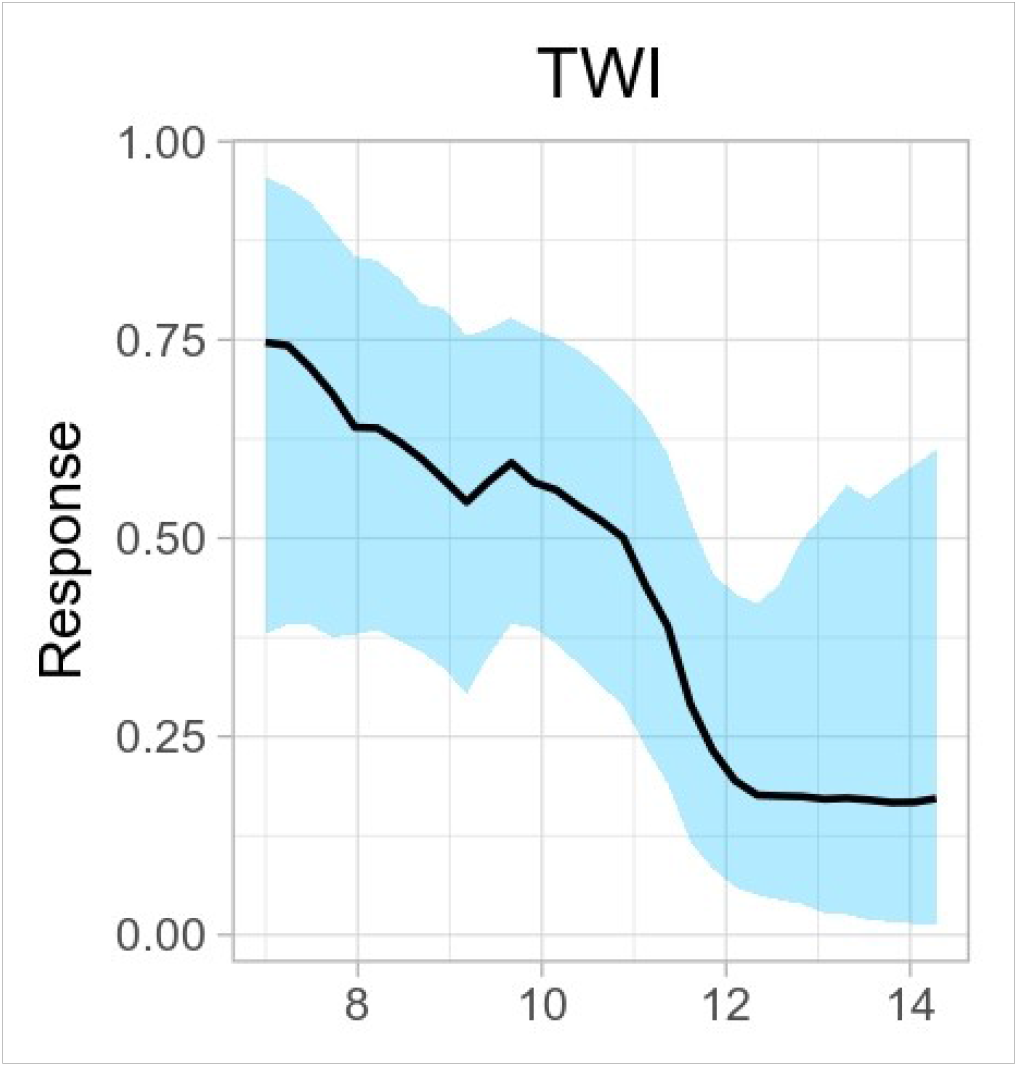
Partial dependence plot for topographic wetness index (TWI)

Finally, terrain roughness (‘tri’) provides a description of the terrain profile and surface heterogeneity. Such heterogeneity plays an important role in catchment-related hydrological responses by driving the flow direction, water runoff velocity, water accumulation, and soil moisture (Bogaart, Troch, 2006). Similarly, topographic variation strongly influences the accumulation and heterogeneity of mountain/alpine snow cover (Grünewald, 2013). Together these factors regulate the water availability in soil, directly influence vegetation and thus can be assumed to be essential for shaping the habitat of *V. turgida*, but because of these multiple associations tri may not itself be the driver of the species’ distribution (Bemmels, 2018). In the Ukrainian Carpathians the species appears to prefer areas of medium to high terrain roughness, where projected habitat suitability reaches its highest value (Figure 4).

**Figure 4.**
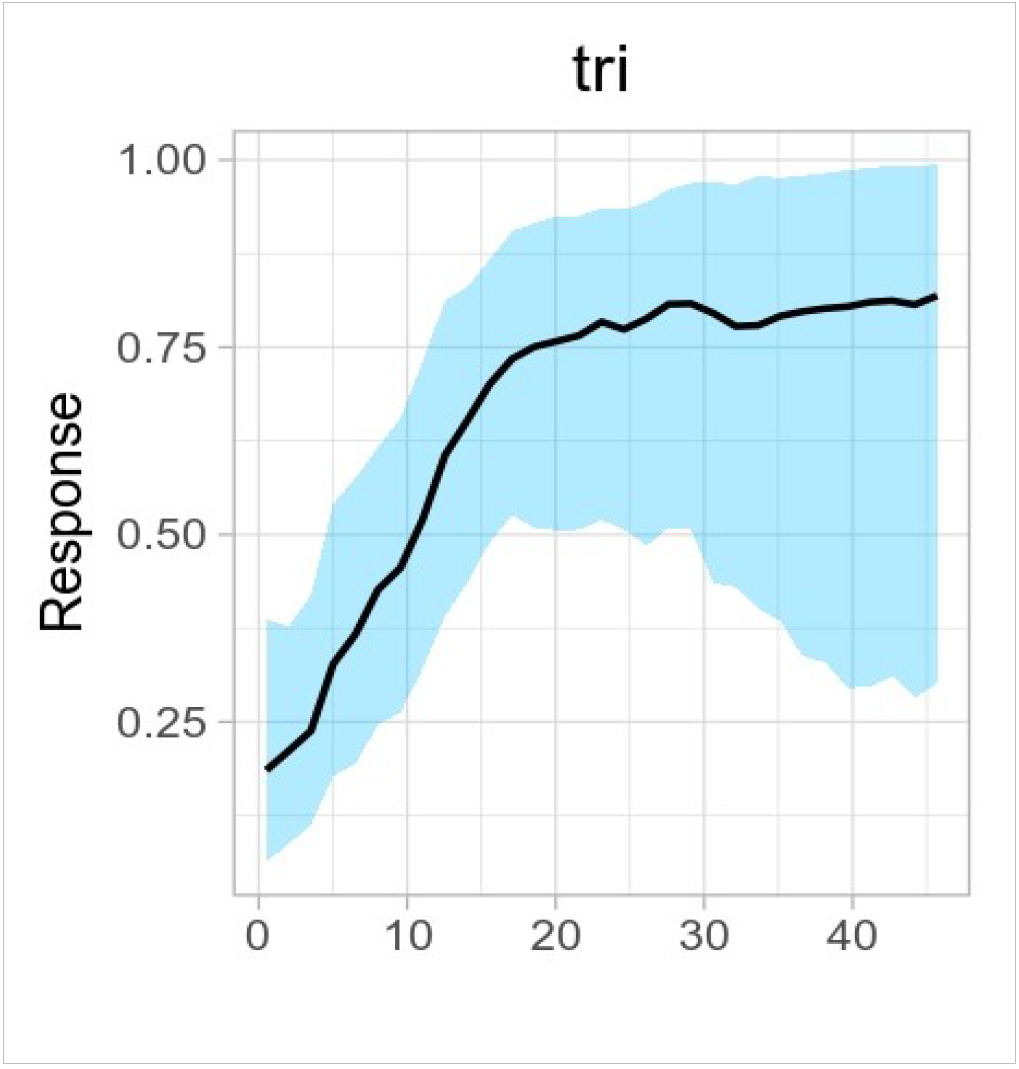
Partial dependence plot for terrain roughness index (tri)

The relationship between shell height and habitat quality as projected by the model was found positive and statistically significant, with larger individuals in areas of preferred topography.

#### 3.2.3 Normalized difference vegetation index (NDVI)

Monthly NDVI were used to build the SDM. There was barely a selection of an informative subset of predictors: most monthly NDVIs were retained for modelling, except for February and June. Accuracy measures for the model are presented in Table 5.

**Table 5.** Accuracy measures for the BART model based on the NDVI

| AUC | TSS | Estimated $\beta \pm SE$ | t-test | p | R<sup>2</sup> | AIC |
| --- | --- | --- | --- | --- | --- | --- |
| 0.894 | 0.667 | 4.790±0.822 | 5.82 | <0.001 | 0.106 | 2996.6 |

Amongst the monthly NDVIs, relatively more important appear those characterizing April and May, when vegetation activity, lower in the winter months, significantly increases (Páscoa et al., 2018).

The relationship between shell height and habitat quality as projected by the model based on monthly NDVIs was found positive and statistically significant, although fairly weak

#### 3.2.4 Soil properties

The following topsoil (0–5 cm) physical and chemical properties were tested: bulk density (cg/cm^3^), clay content (g/kg), coarse fragments (g/kg), sand content (g/kg), silt content (g/kg), cation exchange capacity at pH=7 (mmol(c)/kg), soil organic carbon (dg/kg), pH in water (pH*10), and one derived property, organic carbon density (g/dm^3^), was included. Similar to NDVI, there was a wide selection of predictors used to build the BART model: bulk density, clay content, coarse fragments, silt content, soil organic carbon, and pH in water. Accuracy measures for the model are presented in Table 6.

**Table 6.** Accuracy measures for the BART model based on soil properties

| AUC | TSS | Estimated $\beta \pm SE$ | t-test | p | R<sup>2</sup> | AIC |
| --- | --- | --- | --- | --- | --- | --- |
| 0.914 | 0.640 | 4.920±0.984 | 5.00 | <0.001 | 0.101 | 2766.2 |

Expectedly, pH has been distinguished amongst the selected soil features as the most influential variable. According to the response, higher values of projected habitat suitability are maintained up to an estimated pH of 5.85, after which there is a steady decline (Figure 5), meaning a preference in *V. turgida* towards acidity. In terms of variable importance, next and close to pH is the soil silt content, which reveals a comparable trend: higher values of projected suitability are maintained in habitats where soils contain lesser amounts of silt (we estimate below the level of 47 g/kg); above this estimate projected habitat suitability shows a steady decline.

**Figure 5.**
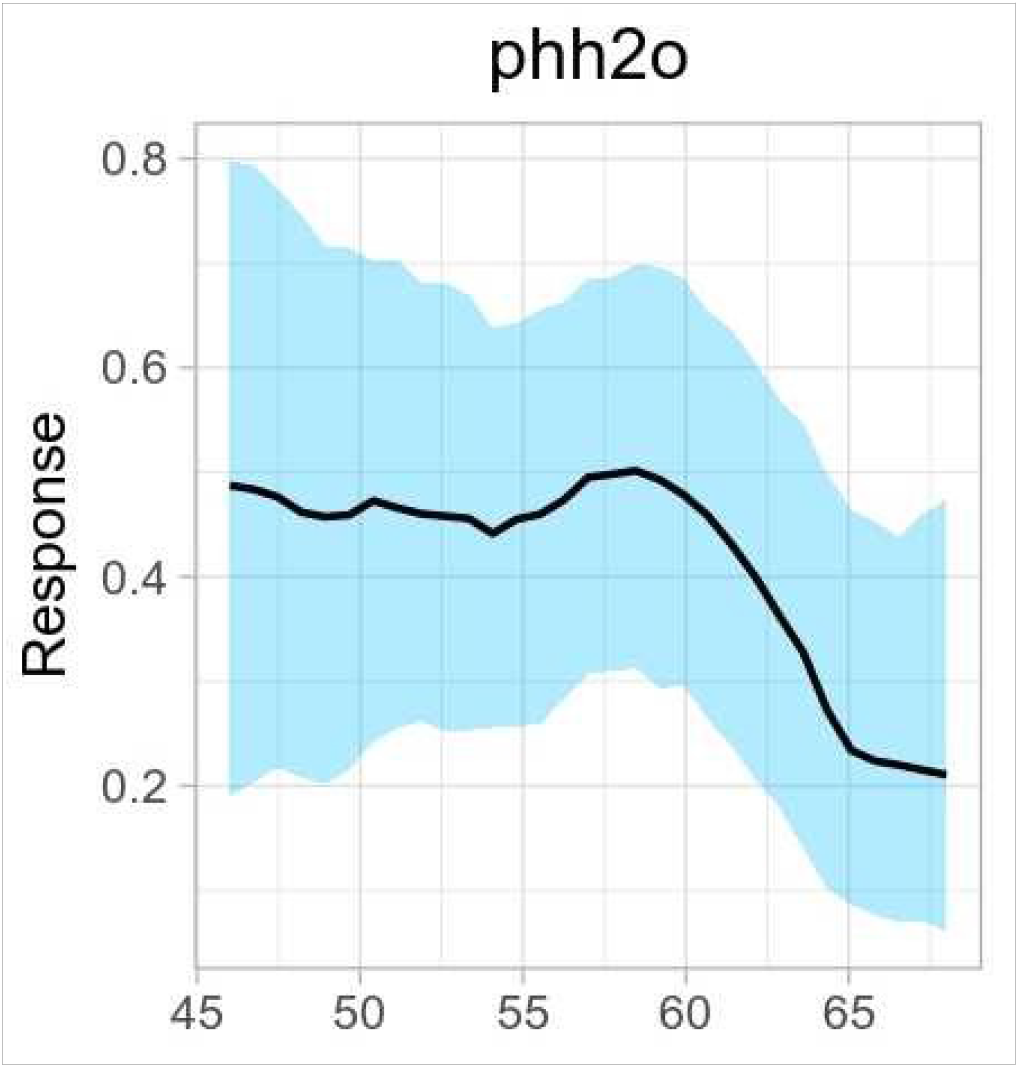
Partial dependence plot for pH water (phh2o)

The relationship between shell height and habitat quality as projected by the model based on soil properties was found positive and statistically significant, although, as in the NDVI case, fairly weak.

Our results are basically in agreement with those of several studies confirming the importance of a number of soil characteristics as determinants of terrestrial gastropod distribution (summarized in: Ondina et al., 2004). General conclusions have been made on the influence of soil properties, which are considered to reflect above all soil acidity and basicity, and secondly soil texture (Ondina et al., 2004).

**Figure 6.**
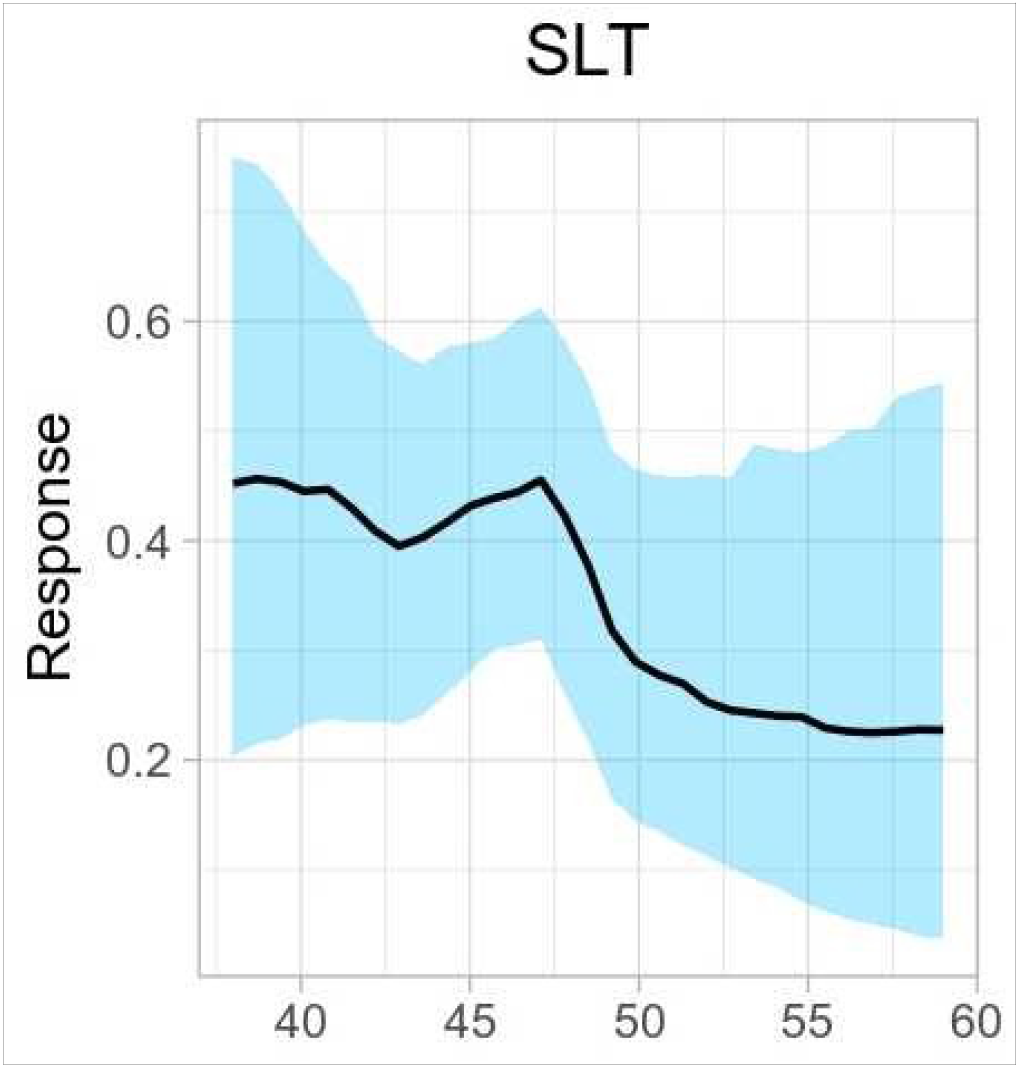
Partial dependence plot for silt content (SLT)

On these grounds the quoted authors (Ondina et al., *op. cit*.) proposed a useful, as regards gastropod distribution, classification of soils based on chemical and physical criteria, where one of the major chemical criteria is pH (consequently, acid and less acid soils), and major physical criteria are textual factors, soil aeration and soil moisture content. Physical criteria allow to distinguish two categories, namely well-drained coarse-textured soils (high proportions of gravel and sand, high aeration, low proportions of silt and clay, low soil moisture content) and wet fine-textured soils (higher proportions of silt and clay, higher soil moisture content). Regarding *V. turgida*, we can say the species prefers acid soils and well-drained coarse-textured soils, for which silt content has served an efficient proxy.

#### 3.2.1 The ENVIREM data set

All 16 climatic and two topographic variables from the ENVIREM dataset were used to produce the BART models. The final recommended variable list consists of three variables: PETColdestQuarter = mean monthly PET of coldest quarter, PETseasonality = monthly variability in potential evapotranspiration, and ‘tri’ = terrain roughness index. On-topic accuracy measures for the model are presented in Table 7.

**Table 7.** Accuracy measures for the BART model based on variables of the ENVIREM dataset

| AUC | TSS | Estimated<br>$\beta \pm \text{SE}$ | t-test | p | R<sup>2</sup> | AIC |
| --- | --- | --- | --- | --- | --- | --- |
| 0.912 | 0.717 | 8.130±0.825 | 9.85 | <0.001 | 0.327 | 2959.1 |

The automated variable selection procedure has put ahead of others ‘tri’, the terrain roughness index, which behaves exactly in the same way as when included to the set of topographic variables (**3.2.2**).

The next two are related to potential evapotranspiration, which is considered to model relationships between water-energy requirements and productivity (Currie, 1991; Field et al., 2005; Fick, Hijmans, 2017). In the first place mean monthly PET of coldest quarter was selected, in the second – monthly variability in potential evapotranspiration. The latter is often viewed as a measure of seasonality of moisture available for vegetation (Zomer et al., 2014) and also is considered to express continentality of the climate (Metzger et al., 2013). In both cases the relationship between the projected habitat suitability values and values of the corresponding variables are hump-shaped, meaning, according to Shelford’s law of tolerance, there is an optimum below or above which a species cannot survive. Tolerance limits regarding the PETColdestQuarter factor could be related to dormancy, whereas PETseasonality could be a reflection of adaptation to the seasonal amplitude in ambient temperature, where differences, either big or small, between seasonal temperature extremes are suggested to be limiting factors.

The relationship between shell height and habitat quality as projected by the model based on the ENVIREM data set was found positive and statistically significant, and appeared notably strong.

## 4. Discussion & Conclusions

Our model species, the terrestrial snail *V. turgida*, provided an opportunity to test hypotheses concerning SDM predictions produced by a number of algorithms commonly used or recently arising due to continuing efforts being put into the refinement of modeling approaches and construction of SDMs (Melo-Merino et al., 2020). Because there is no single ‘best’ algorithm we, as recommended (Qiao et al., 2015), have tested a suite of algorithms for predictive ability and based on the results of these tests selected an algorithm for our particular purpose, which is to describe the environmental niche of the considered species in a variety of perspectives.

In modeling exercises, not only the selection of appropriate modeling techniques, but methods of measuring accuracy are crucial to the outcome (Shabani et al., 2018). Commonly for this purpose diagnostic metrics are used, such as AUC and TSS. However, a high model fit does not necessarily translate into highly consistent spatial or environmental niche predictions (Aguirre-Gutiérrez et al., 2013), and conclusions have been made of the inability of current evaluation metrics to assess the biological significance of SDMs (Fourcade et al., 2018). Indeed, there has been insufficient attention to evaluating the biological meaning of SDM outputs (Wittmann et al., 2016). In our study we have made an attempt to confront the output of the produced SDMs with biological performance data, namely body size of the snails. In general, body size strongly correlates with development times, fecundity, physiological performance, competitiveness and vulnerability to predation, and therefore is considered a fundamental species trait (Wardhaugh et al., 2013). Within species, large individuals often achieve higher reproductive fitness and have greater environmental tolerances than smaller individuals (Shine, 1989). Predictions from SDMs are generally thought to be good indicators of habitat suitability, and thus of species’ performance (Thuiller et al., 2010), consequently SDMs can be validated by checking whether the areas projected to have the greatest habitat quality are occupied by individuals or populations with higher than average fitness (Mammola et al., 2019), and such correlations already have been found (for instance, Thuiller et al., 2010; Nagaraju et al., 2013; Wittmann et al., 2016; Mammola et al., 2019).

We too, hypothesized a positive and statistically significant relationship between observed in the field body size of the snail *V. turgida* and modelled habitat suitability, tested this relationship with linear mixed models, and found that indeed, larger individuals tend to occupy high-quality areas, as predicted by the SDMs. However, by testing several SDM algorithms, we found that some of them performed better, others not so good, in terms of expounding this correlation. In other words, their functional accuracy (Warren et al., 2020) was different. Therefore, marginal R^2^, expressing the variance explained by the fixed terms in the regression models, was adopted as a measure of functional accuracy, and used to rank the SDMs accordingly. In this respect, the Bayesian additive regression trees (BART) algorithm (Carlson, 2020) gave the best result, despite the low AUC and TSS. Interestingly, by functional accuracy the BIOCLIM model outperformed even some machine learning SDM methods.

Our study confirms the possibility to correlate SDM projections with functional traits that serve as proxies for fitness and we propose to use marginal R^2^ to validate these correlations and their strenght.

By restricting our analysis to the BART algorithm only, a variety of sets of environmental variables commonly or less used in the construction of SDMs were explored and tested according to their functional accuracy. In this respect, the SDM produced using the ENVIREM data set gave the best result. Indeed, variables in this data set are worth consideration in SDM applications, especially as many of the variables have direct links to processes important for species ecology (Title, Bemmels, 2018), particularly those related to potential evapotranspiration (PET). However, despite this importance, PET up to now is poorly represented in species distribution modelling (Bradie, Leung, 2017). Satisfactory results were obtained using the sets of topographic and bioclimatic variables, despite reservations against the use of elevation as a predictor or that correlations between climate and species’ distributions could be reflecting the spatial structure of climate rather than real biological process (Beale et al., 2008; etc.). On the contrary, models using vegetation indices and edaphic variables in terms of functional accuracy performed poorly, although the corresponding values of AUC and TSS, considered ‘good’ and ‘very good’, indicate the opposite. We assume the low functional significance of these SDMs is due to scale, because model quality depends not only on the algorithm and applied measure of model fit, but also the scale at which it is used. There are many indications that climate impacts on species distributions are most apparent at macroscales (Vicente et al., 2014), whereas plant biomass or soil may be a more important at the local scale. This also highlights the need to consider an appropriate scale for predictions, as vegetation and edaphic complexity is likely to be degraded by the use of coarse resolution rasters.

Despite some shortcomings, the use of SDMs has allowed to identify some of the important environmental and climatic features that characterize the niche of *V. turgida*. Including other biologically relevant parameters and non-climate variables at apprpropriate scales should contribute important information and help to gain a deeper insight into the niche of the species.

